# Serotonergic Modulation of Spinal Circuitry Restores Motor Function after Chronic Spinal Cord Injury

**DOI:** 10.1101/2022.10.04.510858

**Authors:** Sarita Walvekar, Robert B. Robinson, Hailey M. Chadwick, Rebecca M. Burch, Hanzhang Ding, Steve I. Perlmutter, Samira Moorjani

## Abstract

Electrical stimulation of the nervous system has been employed to enhance the recovery of motor function produced by use-dependent rehabilitation, which is the current gold standard of treatment, following spinal cord injury. However, the therapeutic effects almost always rely on the sustained activation of muscles or neurons, making the benefits largely contingent on continued delivery of stimulation. In the present study, we describe a neuromodulatory intervention that combined intraspinal delivery of serotonergic agonists with use-dependent rehabilitation to restore motor function after a chronic moderate-to-severe cervical contusion in rats that produces impairments in upper-limb movements and dexterity. We show that targeted delivery of quipazine, a broad-spectrum serotonergic agonist, caudal to the lesion increased the effectiveness of physical rehabilitation, leading to substantially improved motor-recovery outcomes in severely-injured, but not moderately-injured, animals. Delivery of quipazine significantly augmented recovery of skilled reach and grasp movements after a severe injury, but moderately-injured animals received no additional benefit from quipazine over physical rehabilitation alone. This difference was perhaps due to a greater loss of serotonin after a severe injury and a resulting environment in which exogenously-applied serotonin can improve circuit function. Our experiments highlight an important role for serotonin in restoration of motor function that is dependent on the severity of the spinal cord injury. They also allude to a potential role for residual serotonin as a biomarker of injury severity. Remarkably, quipazine-mediated behavioral improvements persisted for weeks after termination of neuromodulator delivery, signaling repair of severely-damaged adult spinal circuitry that drives lasting motor recovery.

**Significance Statement:** We describe a neuromodulatory intervention that combined intraspinal delivery of serotonergic agonists with use-dependent physical rehabilitation, which is the current standard of treatment, to promote motor recovery after a chronic moderate-to-severe spinal-contusion injury. Our results show that targeted delivery of serotonergic agonists caudal to the lesion increased the effectiveness of use-dependent rehabilitation, leading to substantially improved motor-recovery outcomes in severely-injured, but not moderately-injured, animals. Notably, therapeutic gains persisted for weeks after termination of neuromodulator delivery—a finding that is both unique and clinically relevant—signaling plasticity induction and repair in chronically-damaged adult spinal circuitry. Our experiments provide important insights into serotonergic modulation of spinal circuitry and highlight a potential role for residual serotonin as a neurochemical biomarker of injury severity.

## Introduction

An estimated 27 million people worldwide live with spinal cord injury (SCI; 1), which often results from trauma suffered during motor-vehicle accidents, falls, or sporting and recreational injuries (2). Among developed regions, annual incidence of traumatic SCI varies from 39 cases per million of the population in North America to 15 – 16 cases per million in Western Europe and Australia (3). SCI can lead to motor deficits in the upper and lower extremities as well as sexual, bowel and bladder dysfunction, depending on the location and severity of the injury (2, 4). Motor deficits after SCI range from weakness in the limbs, abnormal muscle tone and posture, disruption in movement synergies, spasticity, and loss of balance and coordination to complete paralysis (5, 6). Secondary health complications often develop over time due to low levels or the complete lack of movement, further compromising autonomy and quality of life (7). Most spontaneous recovery is limited to the first few months after injury (2, 4). Additionally, a limited number of therapies have been developed for acute SCI (8), but they offer vastly diminished to no benefits to patients with chronic injuries, creating a need for interventions that target chronic motor damage.

Due to the limited intrinsic capacity of the adult central nervous system for repair, existing approaches to motor recovery after chronic injuries seek to leverage spared circuitry for reinforcing functionally-relevant neuronal firing patterns through targeted physical rehabilitation (9). Use-dependent movement interventions, such as robot-assisted body-weight-supported treadmill training (10-12) and constraint-induced therapy (13-15), have been widely explored as treatment options in this effort, but they typically—and at best—lead to marginal recovery of motor function. Greater success has been achieved through hybrid approaches that have combined use-dependent rehabilitation with epidural electrical stimulation of the spinal cord (16-27). However, therapeutic gains achieved using these approaches largely rely on the continued delivery of stimulation. In a small number of studies, residual effects were present (16, 20), but such benefits were appreciably smaller compared to those obtained in the presence of stimulation. This lack of endurance in (the extent of) therapeutic benefits has created a need for development of alternate strategies that produce more lasting changes through modification of the damaged circuitry. Towards this effort, we describe a neuromodulatory intervention that relies on intraspinal delivery of serotonergic agonists below the lesion to restore motor function after chronic SCI.

Serotonin (or 5-hydroxytryptamine; 5-HT) is an important neuromodulator in the spinal cord (28). Serotonergic regulation of spinal-neuron excitability is critical for normal sensorimotor integration and movement production (29). The activation of serotonergic receptors promotes a general increase in motoneuron excitability through amplification of synaptic inputs, depolarization, and increase in firing rates (30). The excitatory effects of serotonin depend on the identity, location and level of receptor activation (30-32). Therefore, serotonergic modulation of spinal circuitry provides a potential avenue for reinstating the (disrupted) excitation–inhibition balance after SCI to promote motor recovery (33, 34).

Previous studies have shown that serotonergic therapies can promote motor gains after SCI (35-43). Delivery of serotonergic-receptor agonists induced stepping and rhythmogenesis in spinalized rats after thoracic injuries through activation of spinal locomotor networks (35-40, 43). Administration of 5-hydroxytryptophan, which is a 5-HT precursor, has also been shown to evoke activity in phrenic motoneurons following a C1 – C2 spinal transection (41). Additionally, serotonergic agonists produced recovery of skilled reach and grasp movements after a C4 injury (42). However, therapeutic benefits did not persist after withdrawal of the neuromodulators in the above studies.

We describe a hybrid intervention that combined intraspinal delivery of quipazine, a non-specific 5-HT_2_ receptor agonist (36, 38, 44), with targeted physical rehabilitation to promote motor recovery in an adult rodent contusion model of chronic cervical SCI. The injury predominantly damages the corticospinal tract, which is critical for control of voluntary upper-limb movements and dexterity (45). Severely-injured rats that received the hybrid intervention showed substantially improved motor-recovery outcomes compared to rats that received physical rehabilitation alone. Importantly, quipazine-mediated benefits outlasted neuromodulator delivery, signaling modification of damaged spinal circuitry that drives lasting improvements in motor function.

## Results

### Experimental design

Experiments were conducted in 20 adult female Long-Evans rats. Rats were trained on a standard forelimb reach-and-grasp behavior (46), using operant-conditioning protocols (47), for food-pellet rewards. Briefly, animals were placed inside clear plexiglass arenas and trained to reach out through a slit in the arena, grasp a food pellet placed on a block outside the arena with their dominant forepaw, and bring the pellet through the slit back into the arena and then to their mouth. Rats were assigned a reach-and-grasp performance score of 1 (or 100%) for successful completion of all the steps in the forelimb-motor sequence. If the rat was unable to touch the pellet, due to inaccuracy in aim or the forelimb not extended far enough, it was counted as a “miss” and was assigned a score of 0. Partial scores were assigned for completion of some—but not all—steps in the motor sequence. To assign partial scores, the motor task was divided into ‘reach’ and ‘grasp,’ with successful completion of the individual components leading to a score of 0.50 on each component (and a combined score of 1). Scoring on reach was binary with rats receiving either 0 or 0.50, whereas scoring of grasp occurred on a quaternary scale (with rats receiving 0, 0.17, 0.33 or 0.50). For example, if the rat dropped the food pellet inside the arena while bringing it to the mouth, it received a reach score of 0.50 and a grasp score of 0.33 (bringing the combined reach-and-grasp score to 0.83). If the pellet, however, was dropped outside the arena, the rat received a reach score of 0.50 and a (lower) grasp score of 0.17 (bringing the total to 0.67). Lastly, if the rat was only able to touch the pellet (but not grasp it), it received a reach score of 0.50 and a grasp score of 0 (making the combined score 0.50). Additional details of the forelimb-motor task and the scoring system can be found in Materials and Methods and Table S1.

Behavioral training on the forelimb-motor task occurred 10X/week, with the sessions divided across the five weekdays. Each behavioral session, in turn, consisted of 20 pellet-retrieval trials, which were preceded by a short warm-up. When rats had attained an average weekly reach-and-grasp performance of 80% or higher on the forelimb-motor task (which typically took 1 – 4 weeks), they received a unilateral (incomplete) moderate-to-severe C4 – C5 spinal-contusion injury on the dominant-forelimb side (48). This type of injury produces marked deficits in extension of the elbow, wrist and digits, consequently, resulting in impaired performance on the previously-mastered skilled reach-and-grasp behavior (49). During week 4 after SCI, rats were implanted with a biocompatible plastic cannula caudal to the lesion in the C7 spinal segment. The cannula was coupled upstream to a pre-programmed battery-operated microinfusion pump (50), which was implanted subcutaneously. Seven weeks after SCI, a pre-therapy assessment on the same task was performed to quantify the functional deficits that remained after spontaneous recovery from the cervical-contusion injury. Therapy was initiated during week 8 after injury, which represents a common time point for a chronic SCI model in rats (51, 52), and was administered during weeks 8 – 13. Therapy consisted of targeted physical rehabilitation combined with intraspinal delivery of quipazine or vehicle. Physical rehabilitation consisted of ten sessions per week of retraining of the impaired forelimb on the pellet-retrieval task and occurred during periods of ongoing neuromodulator or vehicle delivery. To eliminate biases in scoring, all retraining was performed by researchers blinded to the group assignments of the rats. Quipazine or vehicle was delivered intraspinally below the lesion for approximately 4.5 hours/day for a total of six weeks. After completion of therapy, rats continued to be tested in the forelimb-motor task for an additional four weeks to assess retention of motor gains in the absence of the neuromodulator. A terminal surgery was performed after completion of post-therapy follow-up to assess the integrity and location of the cannula. A detailed experimental timeline is shown in Figure 1A.

**Figure 1.**
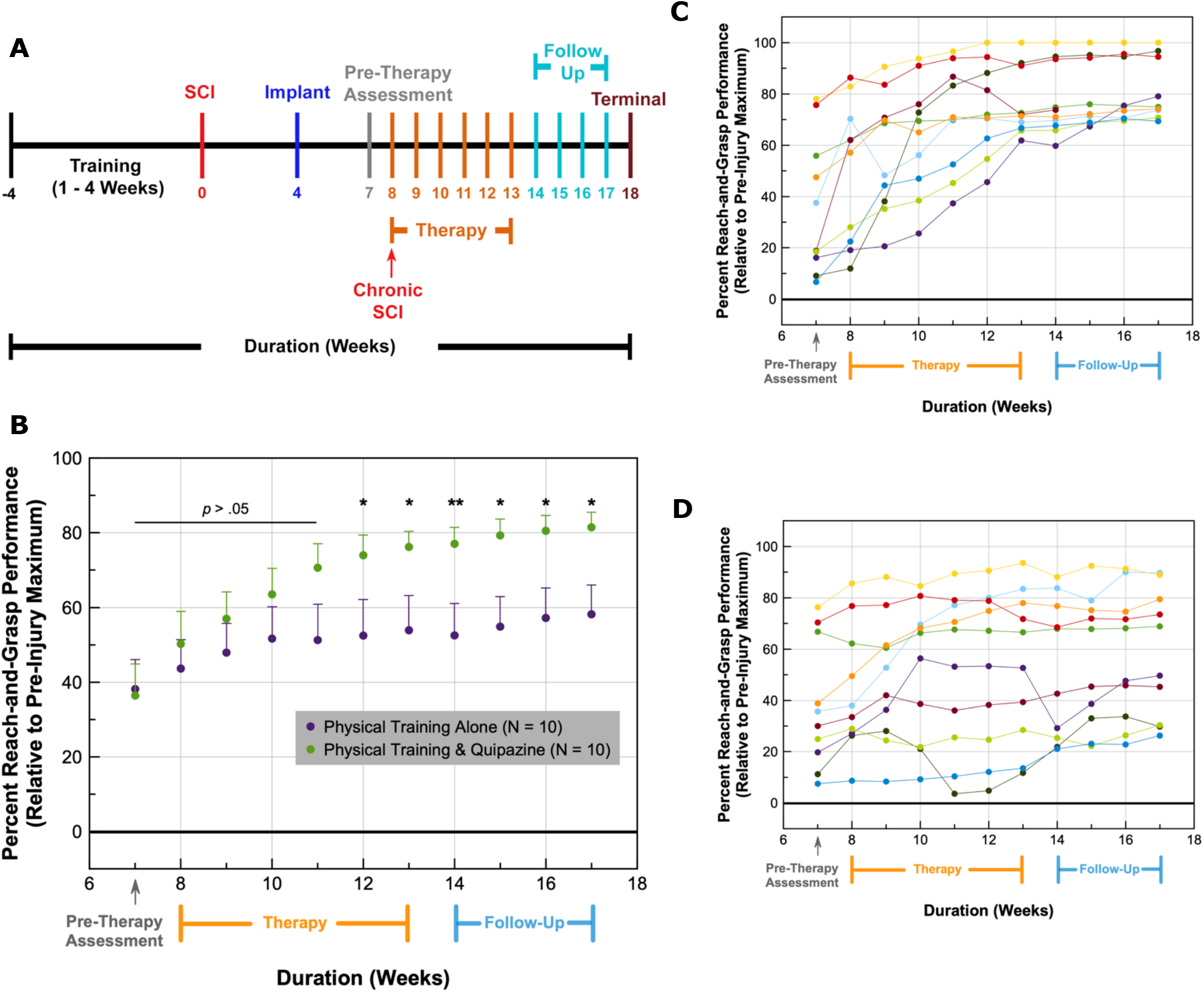
Intraspinal quipazine enhances motor recovery after chronic SCI. (A) Experimental timeline. (B) The reach-and-grasp scores of injured rats, expressed as a percentage of their individual pre-SCI maximum, were used to quantify forelimb-motor performance before, during and after therapy. Data points represent weekly averages of all the rats in their respective intervention groups. ‘N’ indicates the total number of rats in each group. Error bars represent standard error of the mean. Only positive error bars are shown in the plot to reduce clutter. *p* values were obtained using paired-samples *t* tests that compared performance scores of animals in the two intervention groups. \**p* ≤ .05; \*\**p* ≤ .01. (C, D) Individual reach-and-grasp performance profiles of rats, whose means are shown in (B), that received quipazine (C) or vehicle (D) during therapy. Motor-recovery profiles are color-coded in the plots to reflect impairment matching of rats randomly assigned to the two groups.

Note that rats in both intervention groups received retraining on the reach-and-grasp pellet-retrieval task during weeks 7 – 17. Retraining on pellet retrieval served as both targeted physical rehabilitation and a means to quantify behavioral recovery promoted by our interventions.

### Intraspinal quipazine improves motor-recovery outcomes after chronic SCI

Chronically-injured rats, after a moderate-to-severe cervical SCI, received physical rehabilitation combined with intraspinal delivery of (20 µg/week) quipazine or vehicle.

Depending on their pre-therapy (or week 7) reach-and-grasp scores, impairment-matched animals were randomly assigned to the two intervention groups. Impairment matching was used to account for the variability in motor deficits across animals arising from the large range of injury severities tested in our study (see Materials and Methods and Table S2 for additional details). Paired-samples *t* tests were employed for statistical comparisons between impairment-matched (case–control) groups (53, 54). Figure 1B shows the average weekly reach-and-grasp performance of all rats in the two intervention groups during the pre-therapy, therapy and follow-up periods. The reach-and-grasp performance of the injured rats is reported as a percentage of their individual pre-SCI maximum scores.

Injured rats that received physical training combined with intraspinal delivery of quipazine showed greater improvement in their forelimb-motor performance compared to rats that received physical training alone (Figure 1B). During the pre-therapy assessment, the reach-and-grasp performance of rats in the two intervention groups was indistinguishable from each other (*t*(9) = -0.845, *p* > .05, paired-samples *t* test). Starting with the first week of the intervention, rats that received physical training and quipazine showed greater improvement in their performance scores relative to those that received physical training alone, and this difference in motor performance widened as therapy progressed. The motor-performance scores of rats in the two groups, however, were not statistically different from each other during the first four weeks of the intervention (*i.e.,* weeks 8 – 11). Beginning with the fifth week of the intervention (or week 12 after SCI), rats that received quipazine performed significantly better than those that received physical training alone (and this trend continued into the follow-up period). At the end of the six-week therapy period, the average forelimb-skilled performance of rats that received the combined intervention was 76% ± 4%, while rats that received physical training alone recovered 54% ± 9% of their pre-injury ability. The above changes, as well as those in the following sections, are reported as mean ± standard error (unless otherwise indicated).

Figures 1C and 1D show the individual motor-recovery profiles of rats that received intraspinal quipazine and vehicle, respectively. Recovery profiles of rats that started at similar functional injury levels, as deduced through their week 7 reach-and-grasp scores, are represented by the same color in the two plots to illustrate differences in intervention-mediated recovery between them (*e.g.,* yellow curves depict the profile of the least injured rat in the two groups, while the darker-blue curves represent the most severely injured rat). Because rats in the two intervention groups were impairment-matched, and they spanned a range of impairments, there was a high—but comparable—variability in motor-performance scores across animals within the two groups during the pre-therapy week. Pre-therapy standard deviations were 27% and 25% for rats in the quipazine and vehicle groups, respectively. Variability in motor performance gradually decreased as therapy progressed in rats that received the combined intervention, with the standard deviation dropping to 13% at the end of the six-week therapy period (Figures 1B and 1C). In contrast, variability in the physical-training-alone group continued to remain high, and the standard deviation was 29% at the end of therapy (Figures 1B and 1D). Notably, the difference in variability between the two groups at the end of therapy was primarily due to rats with higher injury severity (or lower pre-therapy reach-and-grasp scores). In the combined intervention group, these rats achieved larger therapeutic gains relative to the moderately-injured animals in the group, which resulted in comparable performance scores across animals and reduced variability over time (Figure 1C). This decreasing trend in performance variability was not seen in rats that received physical training alone (Figure 1D).

### Quipazine-mediated gains persist beyond the duration of therapy

We assessed the persistence of quipazine-mediated benefits by assessing the motor performance of injured rats for four additional weeks, after the end of therapy, in the absence of quipazine (Figures 1A, 1B and 1C). During the four-week follow-up period, rats received the same amount of physical training (10X/week), which also served as a means to quantify the behavioral recovery promoted by our interventions. A comparison of the motor-performance scores in the combined intervention group during the follow-up period (*i.e.,* weeks 14 – 17) found significant differences between the four weeks (*X*^2^(3, N = 9) = 15.835, *p* = .001, Friedman test; N, sample size). Note that one of the rats was excluded from the statistical analysis due to listwise deletion from some missing post-therapy scores, reducing the sample size to nine. Next, we compared performance scores between the last week of therapy (*i.e.,* week 13) and the first week of post-therapy follow-up (*i.e.,* week 14) of rats in the quipazine group and found that there were no significant differences in the scores between the two weeks (*t*(9) = 1.764, *p* > .05, paired-samples *t* test; Figure 2). In contrast, significant increases were observed during weeks 15 – 17 compared to scores obtained during week 13 in the combined intervention group (Figure 2). While the increases in scores were significant during weeks 15 – 17, the size of the effect ranged from very small to small (0.19 – 0.37), and this small increase may have been promoted by the continued physical training during the follow-up period. Details of effect-size calculations and thresholds used for qualitative interpretation of the magnitude of observed effects can be found in Materials and Methods and Table S3.

**Figure 2.**
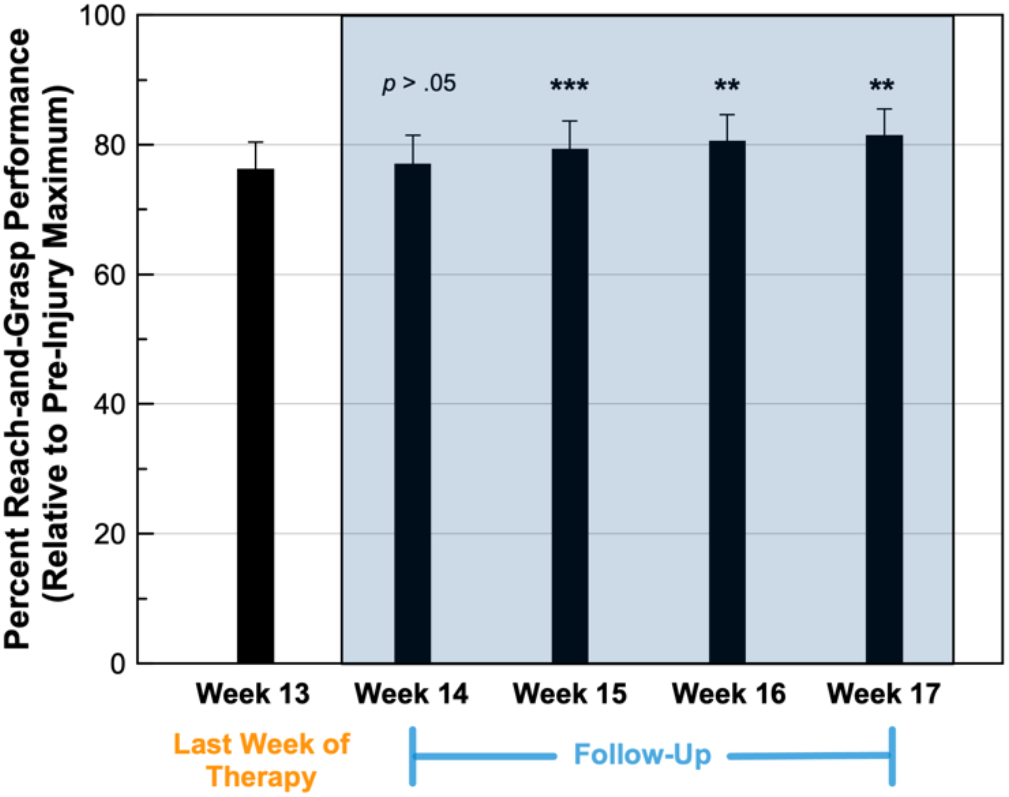
Intraspinal quipazine produces lasting motor recovery after chronic SCI. Bar plot shows the average reach-and-grasp performance during the last week of therapy and the four weeks of post-therapy follow-up (shown in the blue-shaded region) for animals that received physical training combined with intraspinal delivery of quipazine (see Figures 1B and 1C). Error bars represent standard error of the mean. *p* values and asterisks indicate statistical comparisons between the performance of rats during the corresponding follow-up week and week 13. *p* values were obtained using paired-samples *t* test or Wilcoxon signed-rank test, depending on the distribution of the data. \*\**p* ≤ .01; \*\*\**p* ≤ .001.

### Quipazine-mediated benefits are dependent on the injury severity

Figure 3A shows the average therapeutic gains (= mean post-therapy performance – pre-therapy performance) produced by our interventions. The mean post-therapy performance was calculated by averaging the reach-and-grasp scores obtained during weeks 14 – 17. Therapeutic gains were 2.4-fold larger with quipazine (43% ± 7%) compared to physical training alone (18% ± 5%). We used a paired-samples *t* test to compare performance gains between impairment-matched rats, which revealed a significant difference between the two intervention groups (*t*(9) = 2.925, *p* = .02; effect size = 1.57, very large effect).

**Figure 3.**
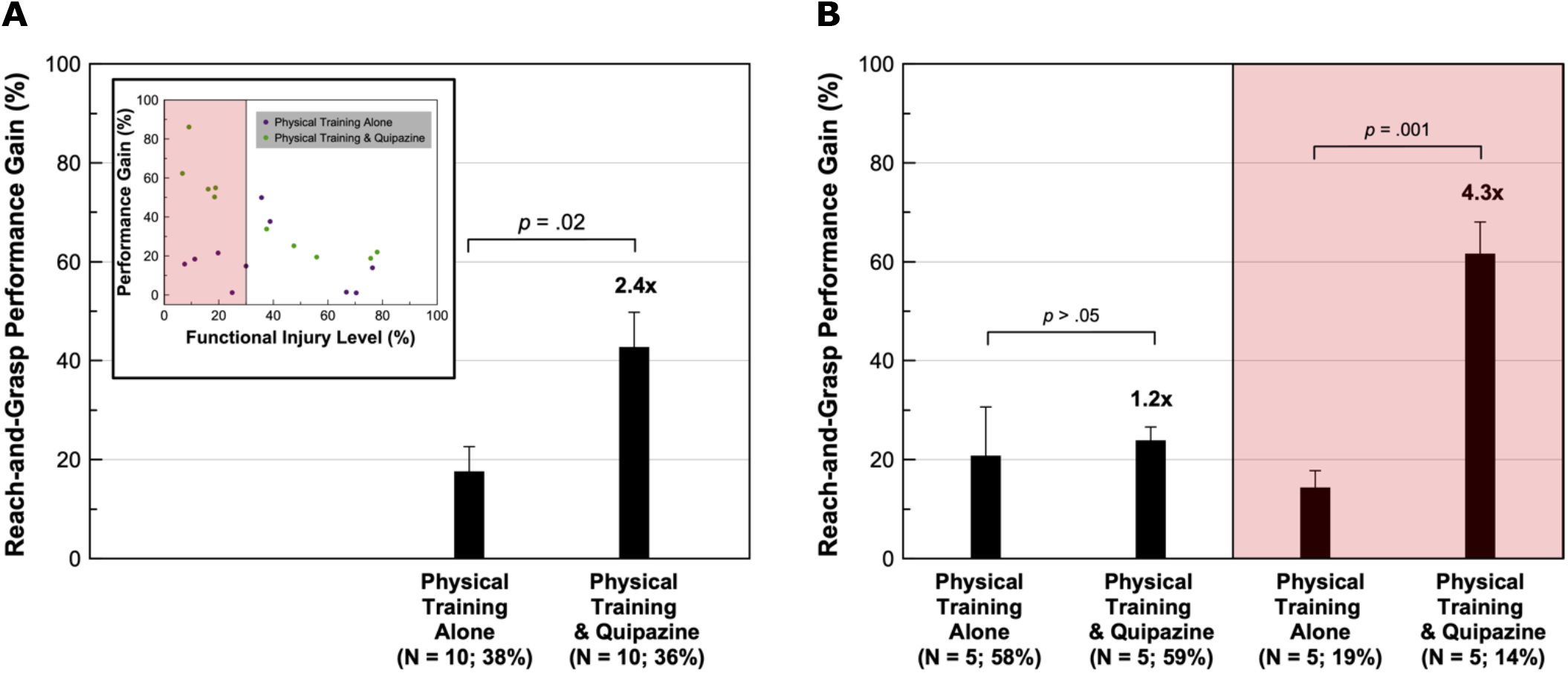
Intraspinal quipazine augments motor gains after severe SCI. (A) Bar plot shows the average reach-and-grasp performance gains for animals that received physical training and intraspinal delivery of vehicle or quipazine. Inset shows the performance gains as a function of injury severity for individual rats in the two intervention groups. (B) Bar plot shows intervention-mediated performance gains after a moderate *vs.* severe SCI. Severely-injured rats are shown in the pink-shaded region, while moderately-injured rats are shown in the unshaded region of the plots in the inset and (B). Numbers (in bold) above the bars quantify the relative performance of rats that received the combined intervention over those that received physical training alone. Error bars represent standard error of the mean. *p* values were obtained using paired-samples *t* tests. Number of rats in each group, along with the average injury severity, are reported in parentheses below the bars.

We also plotted therapeutic gains as a function of the injury severity, quantified by the pre-therapy (or week 7) performance scores, for individual rats (Figure 3A, inset). We found that quipazine-mediated gains were strongly correlated to the injury severity. Rats with higher injury severity (or lower pre-therapy reach-and-grasp scores) attained larger therapeutic gains (Pearson’s correlation coefficient = 0.89). In contrast, therapeutic gains showed a very weak correlation to injury severity in the physical-training-alone group (Pearson’s correlation coefficient = 0.28). Next, we classified rats with pre-therapy performance scores of less than or equal to 30% as severely injured (shown in the pink-shaded region in Figure 3A, inset) and rats whose pre-therapy performance was greater than 30% as moderately injured (shown in the unshaded region in the inset). We found that there was no significant difference in intervention-mediated reach-and-grasp improvements between moderately-injured animals that received quipazine or vehicle (*t*(4) = 0.413, *p* > .05, paired-samples *t* test; Figure 3B, unshaded region). Delivery of quipazine, therefore, produced no additional benefit in animals with a moderate SCI. In stark contrast, the average reach-and-grasp performance gain was 4.3-fold larger in severely-injured rats that received physical training and quipazine compared to those that received physical training alone (*t*(4) = 8.066, *p* = .001, paired-samples *t* test; effect size = 6.05, very large effect; Figure 3B, pink-shaded region). These results indicate that intraspinal quipazine significantly augmented motor gains only after severe SCI.

### Intraspinal quipazine enhances recovery of skilled reach and grasp movements after severe cervical SCI

The forelimb-motor behavior—that served as targeted physical rehabilitation—primarily consists of two coordinated functional movements: reach and grasp (49, 55, 56). These two movements may be differentially affected after a cervical SCI. To determine the effect of quipazine on the recovery of these movements, we compared intervention-mediated changes in the reach and grasp motor performances, assessed individually (see Materials and Methods and Table S1 for description of behavioral scoring). Figure 4A shows the average weekly reach performance of injured rats as a percentage of their individual pre-SCI maximum scores. The average pre-therapy scores of rats in the two intervention groups were similar (57% ± 10% and 53% ± 11% for rats receiving vehicle and quipazine, respectively). Starting with the first week of the intervention, rats that received physical training and quipazine showed greater improvements in their reach ability relative to those that received physical training alone, and this difference widened as therapy progressed. Importantly, performance variability decreased over time in the combined intervention group, with all rats recovering 100% of their pre-injury reach ability at the end of the post-therapy follow-up. In comparison, recovery of reach in the impairment-matched rats that received physical training alone ranged between 46% to 100%.

**Figure 4.**
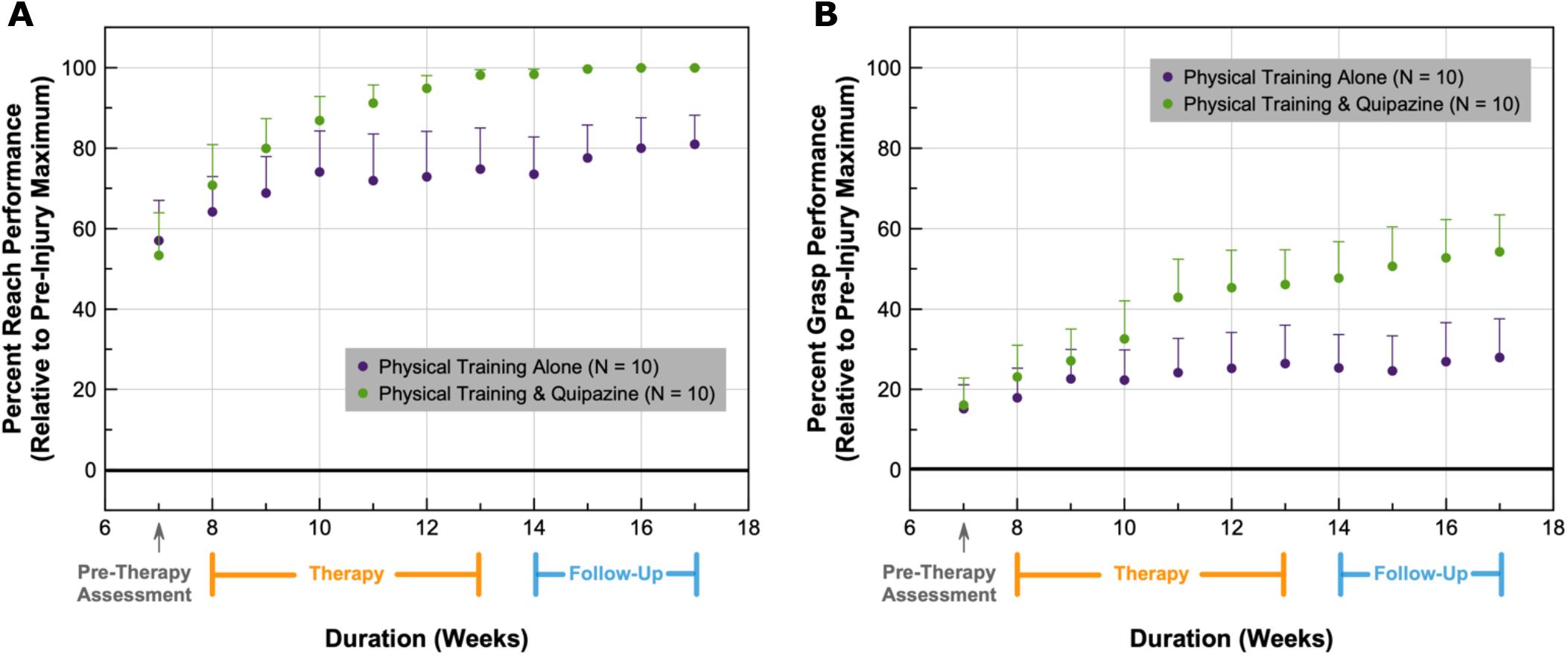
Intraspinal quipazine improves recovery of skilled reach and grasp movements after chronic cervical SCI. (A, B) The average weekly reach (A) and grasp (B) performance of rats, reported as a percentage of their individual pre-SCI maximum scores, is plotted for the pre-therapy, therapy and follow-up periods. Rats received physical training and intraspinal delivery of vehicle or quipazine during the six-week therapy period. Error bars represent standard error of the mean.

In comparison to deficits in reach, grasping ability was more impaired after the cervical contusion with rats starting at appreciably lower pre-therapy (grasp) baselines: 15% ± 6% and 16% ± 7% for rats receiving vehicle and quipazine, respectively (Figure 4B). Although trends in grasp performance were similar to those of reach (*cf.* Figures 4A and 4B), gains were smaller, with rats recovering 54% of their pre-injury grasp ability at the end of the follow-up period in the presence of quipazine and only 28% with physical training alone. Finally, in contrast to improvements in reach, performance variability in recovery of grasp continued to remain high during the therapy and follow-up periods and was comparable between the two intervention groups.

Figure 5 shows the average therapeutic gains in reach and grasp performances produced by our interventions. The average gain in reaching ability was 2.2-fold larger in injured rats that received physical training and quipazine compared to those that received physical training alone (*t*(9) = 2.642, *p* = .03, paired-samples *t* test; effect size = 0.96, large effect). Quipazine-driven gains in grasp were even larger with injured rats recovering 3.1-fold more grasp compared to those that received physical training alone (*t*(9) = 2.622, *p* = .03, paired-samples *t* test; effect size = 1.07, large effect). These results indicate that intraspinal delivery of quipazine significantly improved recovery of both skilled reach and grasp movements after chronic cervical SCI.

**Figure 5.**
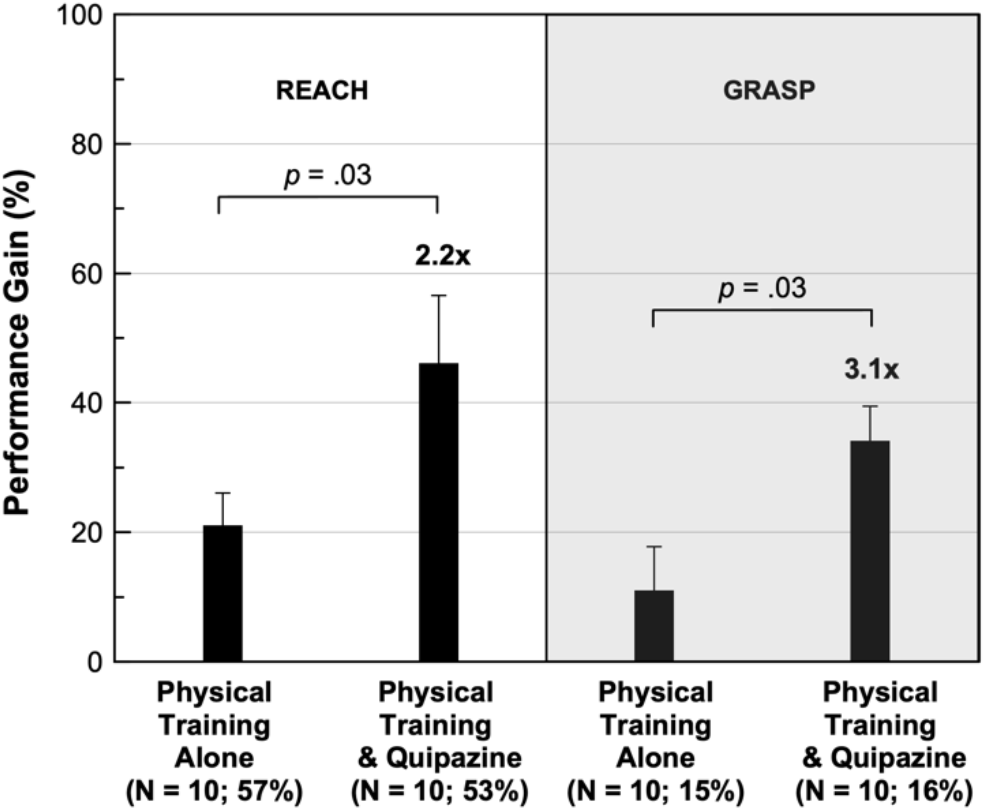
Intraspinal quipazine promotes greater recovery of skilled movements after chronic SCI. The reach (left unshaded panel) and grasp (right gray panel) performance gains are plotted for injured rats that received physical training and intraspinal delivery of vehicle or quipazine. Numbers (in bold) above the bars quantify the performance of rats that received the combined intervention relative to those that received physical training alone. Error bars represent standard error of the mean. *p* values were obtained using paired-samples *t* tests. Number of rats in each group, along with the average pre-therapy reach/grasp baseline, are reported in parentheses below the bars.

Finally, we plotted the performance gains in reach and grasp as a function of injury severity, using the injury-severity classification developed in inset of Figure 3A, to characterize the effect of our interventions after a moderate *vs.* severe SCI. Using paired-samples *t* tests, we found that there were no significant differences in the reach or grasp gains produced by our interventions in the moderately-injured animals (*t*(4) = -0.320, *p* > .05 and *t*(4) = 0.784, *p* > .05 for reach and grasp, respectively; Figure 6A). In striking contrast, delivery of quipazine, caudal to the lesion, produced substantially larger improvements in both reach and grasp ability after severe cervical SCI (Figure 6B). Severely-injured rats that received quipazine showed a 3.1-fold larger gain in their reach ability relative to the physical-training-alone group (*t*(4) = 8.139, *p* = .001, paired-samples *t* test; effect size = 4.01, very large effect). Quipazine-mediated gains in grasp after severe SCI were also larger (*t*(4) = 3.523, *p* = .02, paired-samples *t* test; effect size = 10.89, very large effect). Individual gains in grasp were less than 3% in the absence of quipazine after severe SCI. In the presence of quipazine, the average gain in grasp in the severely-injured animals was 36%. These results demonstrate the effectiveness of quipazine in promoting recovery of skilled reach and grasp movements after a severe, but not moderate, SCI.

**Figure 6.**
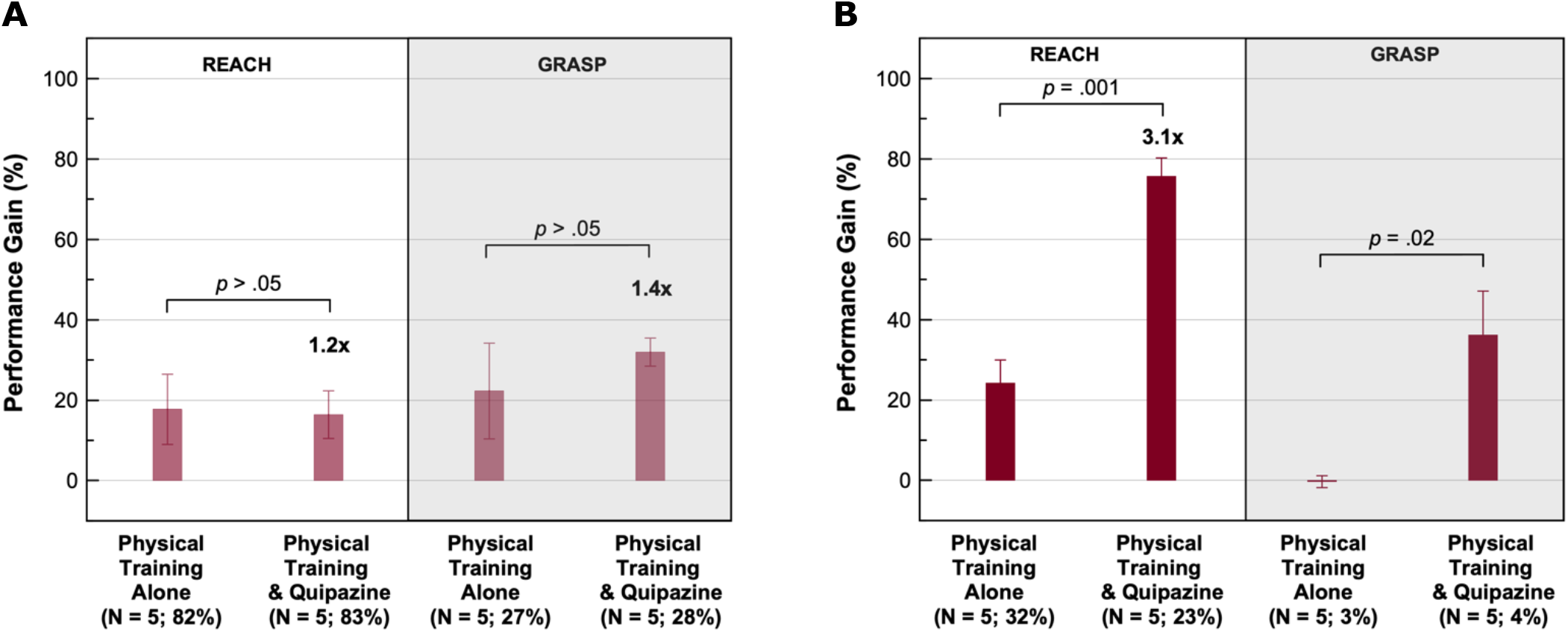
Intervention-mediated forelimb-performance gains after a moderate *vs.* severe SCI. (A, B) The reach (left panel) and grasp (right panel) performance gains after a moderate (A) *vs.* severe (B) SCI are plotted for rats that received physical training and intraspinal delivery of vehicle or quipazine. Numbers (in bold) above the bars quantify the performance of rats that received the combined intervention relative to the physical-training-alone group. Error bars represent standard error of the mean. *p* values were obtained using paired-samples *t* tests. Number of rats in each group, along with the average pre-therapy reach/grasp baseline of the group, are reported in parentheses below the bars. Note that quipazine-mediated gains in grasp are not reported in the severely-injured cohort (B, right panel) due to the lack of gains in the physical-training-alone group (which produced an inflated ratio when comparing gains between the two interventions).

## Discussion

This study describes a neuromodulatory intervention that relies on intraspinal delivery of serotonergic agonists to promote motor recovery after a chronic moderate-to-severe cervical SCI. Chronic motor deficits are one of the most debilitating symptoms experienced by individuals with SCI, and restoration of upper-extremity function has been identified as an important recovery priority (57). Impairments in upper-extremity function can compound difficulties in other areas—such as bowel and bladder management, both of which have also been identified as recovery priorities (57)—to further reduce independence and quality of life. Physical rehabilitation and use-dependent movement interventions are the current standard of treatment after SCI, but they offer limited benefits, especially after chronic trauma (10-15). Our results demonstrate that targeted delivery of quipazine, a non-specific 5-HT_2_ agonist (36, 38, 44), caudal to the lesion substantially increased the effectiveness of physical rehabilitation in promoting forelimb-motor recovery in an adult rodent contusion model of chronic severe, but not moderate, cervical SCI. Injured rats in our study received physical rehabilitation combined with intraspinal delivery of quipazine or vehicle. All rats that received quipazine recovered 100% of their pre-injury reach ability and showed substantial improvements in grasp, which led to sizable gains in forelimb-motor performance and reduced post-therapy performance variability in the combined intervention group. In comparison, physical rehabilitation alone provided significantly reduced benefits, which were also more variable across animals. This difference was due to the beneficial effect of quipazine on motor recovery after severe SCI. While recovery of skilled reach and grasp movements was comparable between the two intervention groups in moderately-injured animals, quipazine significantly augmented forelimb-motor gains after severe SCI. Gains in reach were substantially smaller in the absence of quipazine after severe SCI, and physical training alone was completely ineffective in the rescue of skilled grasping movements in severely-injured animals. Our data, therefore, highlight an important role for serotonin in restoration of motor function after chronic SCI that is dependent on the severity of the injury. Remarkably, quipazine-driven improvements in function outlasted neuromodulator delivery, suggesting that our serotonergic intervention had modified the underlying spared, but malfunctional, circuitry to drive lasting motor recovery.

### Potential mechanisms underlying quipazine-driven motor recovery after chronic severe SCI

The dendrites of motoneurons have voltage-dependent channels that can generate strong sodium and calcium persistent inward currents (PICs), which result in the characteristic sustained motor-unit firing pattern. The amplitude of the inward current is proportional to the level of neuromodulatory input from the brainstem, and serotonin plays an important role in the modulation of PICs (58). SCI results in a loss (or reduction) of brainstem-derived serotonin. In the days following SCI (*i.e.,* during the acute phase of the injury), motoneurons have small PICs and reduced excitability, and spinal reflexes are weak or absent, which leads to impairments in motor function or total paralysis (33, 58).

Over time, as the injury enters the chronic phase, motoneurons compensate for the loss of serotonergic input and degeneration of descending (serotonergic) axons through an overexpression of 5-HT_2_ receptors—primarily 5-HT_2A_ (59-61) and 5-HT_2C_ (33, 62) subtypes—caudal to the lesion. Moreover, these receptors switch to a constitutive phenotype (33, 63), which restores the PICs and, in turn, motoneuron excitability to enable sustained muscle contractions (in the continued absence of brainstem-derived serotonin) that, ultimately, lead to some spontaneous recovery of motor function.

Other studies have reported that low levels of serotonin (2 – 12%), derived in part from spinal serotonergic neurons associated with the autonomic system, persist in the spinal cord below a chronic transection (28, 64, 65). This combined with the enhanced sensitivity of motoneurons to residual monoamines after a chronic injury (51, 52, 66, 67) have also been postulated to be critical to the redevelopment of PICs. Bennett and coworkers found that very low levels (0.5 – 3%) of serotonin were sufficient to facilitate spinal reflexes after a chronic SCI relative to the dose required after an acute injury of the cord (51, 52, 67).

Our study found that intraspinal delivery of quipazine, caudal to the lesion, provided no additional benefit over physical rehabilitation alone after a moderate SCI. Considering that we implemented a chronic model of incomplete cervical SCI, it is expected that there would be some brainstem-derived serotonin caudal to the lesion in these animals. It is, therefore, not surprising that the residual serotonin—derived from both spared descending pathways and autonomic sources—combined with the supersensitivity of motoneurons after a chronic injury precluded any advantage of our serotonergic intervention after a moderate SCI. In contrast, delivery of quipazine significantly improved motor-recovery outcomes after a severe SCI in chronically-injured animals. This striking difference may be due to a greater loss of brainstem-derived serotonin after a severe injury and a resulting environment in which exogenously-applied serotonin can improve circuit function.

In the absence of normal inhibitory input, especially from spinal interneurons that are also regulated by descending tracts (68, 69), self-sustained depolarizations, arising from compensatory processes, can produce involuntary muscle spasms or spasticity (33, 58). Spasticity is a debilitating symptom that affects 65 – 78% of individuals with chronic SCI (5), further contributing to motor dysfunction. Since motoneuron function and normal movement production require both excitatory and inhibitory control (58), and serotonergic modulation is expected to increase motoneuron excitability (70), we speculate that the incomplete nature of our cervical injury spared enough inhibitory circuits, and descending tracts that regulate them (68, 69), to allow for restoration of movements without occurrence of appreciable levels of spasticity. While spasticity was not quantitatively assessed in our study, the large extent of motor recovery observed in the severely-injured animals that received quipazine supports this premise.

Lastly, several studies have found that monoamines, primarily serotonin and norepinephrine, increase the sensitivity of motoneurons to both excitation and inhibition (71-73). Furthermore, a wide variety of spinal interneurons are strongly influenced by serotonergic inputs to the cord (74-77) through activation of 5-HT_1B_ and 5-HT_1D_ receptors (77, 78). Therefore, delivery of a non-specific serotonergic agonist, such as quipazine, may also have modulated the inhibitory input to motoneurons and the activity of spinal interneurons (in addition to enhancing motoneuron excitability; 70) to promote motor recovery after a severe and chronic injury of the spinal cord. Larger-scale investigations, incorporating *in vivo* electrophysiology and genetic manipulations, such as transgenic animals and viral vectors, are needed to elucidate how (and which) specific subsets of spinal neurons and serotonergic receptors contribute to the motor recovery mediated by delivery of quipazine caudal to the lesion after chronic severe SCI.

### Therapeutic benefits outlast neuromodulator delivery

Our results show that delivery of quipazine caudal to the lesion substantially improved motor function after chronic severe SCI. Importantly, quipazine-mediated benefits persisted for at least four weeks after the end of therapy (in the absence of quipazine; Figure 2). The maintenance of quipazine-driven recovery for weeks beyond the duration of therapy is both a unique and clinically-relevant finding. It distinguishes our neuromodulatory intervention from most electrical-stimulation approaches whose benefits are largely contingent on the continued delivery of therapy (16-27, 79, 80), and residual effects, if present, are either appreciably smaller (16, 20) or decay within minutes to hours (79, 80).

Serotonergic agonists have been used extensively to promote motor recovery after SCI (35-43). However, only transient benefits were observed in these studies. In one study by Antri and coworkers, daily injections of quipazine and 8-OHDPAT (which is an agonist for 5-HT_1A_ and 5-HT_7_ receptors) for one month, after a T8 – T9 SCI, led to long-term improvements in locomotion that lasted for up to 46 days after the end of neuromodulator administration (81). While the longevity of motor gains is striking, serotonergic therapy was initiated the day after SCI. Serotonergic interventions were, furthermore, delivered during the acute or subacute phase of the injury in all the above studies (35-43, 81). However, the majority of SCI patients have injuries that are in the chronic phase (82). Our study, in which quipazine delivery was initiated eight weeks after SCI (which represents a common time point for a chronic SCI model in rats; 51, 52), provides critical evidence for the important role of serotonin in repair after a chronic and severe contusion of the spinal cord. The substantial and lasting recovery promoted by our serotonergic intervention further underscores its clinical relevance for motor restoration after severe chronic damage.

Lastly, a previous study from our laboratory, by McPherson *et al*., used targeted activity-dependent spinal stimulation to strengthen spared motor pathways after an incomplete cervical SCI (83). Delivery of intraspinal stimulation caudal to the lesion, triggered by residual electromyographic (EMG) activity in the impaired forelimb, produced lasting motor recovery in a rodent contusion model of chronic cervical SCI (similar to the one used in the current study). The average pre-therapy baseline across the three intervention groups in the McPherson *et al*. study was 13% ± 2%, relative to pre-SCI levels, with no between-group differences in performance before the start of treatment. Forelimb-motor performance was calculated using a binary measure that relied only on the number of successes in the pellet-retrieval task (which is different from the metrics employed in the current study). Using the McPherson-*et-al*. metric (for direct comparison), we found that rats in our (current) study were more injured averaging at 4.0% ± 1.9% when all rats were included (N = 20) and 0.2% ± 0.2% in the severely-injured cohort (N = 10) with no significant performance differences between animals in the two intervention groups before the start of therapy (*Z* = -0.943, *p* > .05, Wilcoxon signed-rank test). Moreover, animals with a pre-intervention score of zero did not benefit from activity-dependent stimulation unless they achieved an average of at least one success in either of the first two weeks of therapy, perhaps due to the lack of enough spared pathways necessary for an EMG-based intervention. In contrast, six out of the ten animals that received quipazine had a pre-intervention score of zero, and they all showed substantial and lasting improvements in motor function. These comparisons highlight a critical role for serotonin in long-term motor restoration after chronic severe SCI that is perhaps less reliant on spared pathways and could involve axonal sprouting and new synaptic connections (84, 85).

### Serotonin as a potential neurochemical biomarker of SCI severity

The complexity and heterogeneity of spinal-contusion injuries pose significant barriers to clinical translation of therapies, which often target specific pathways and may only be effective for a narrow range of injury severities and biochemical-damage profiles. Biomarkers are, therefore, needed, so clinicians can track SCI progression, predict recovery outcomes, and use that information to effectively tailor treatments (86, 87). Our results show that quipazine-driven gains were strongly correlated to the severity of the injury (Figure 3A, inset). While our hybrid intervention promoted significant and substantial motor gains after a chronic and severe injury of the cord, moderately-injured animals received no additional benefit from quipazine over physical rehabilitation alone. This strong dependence of therapeutic gains on injury severity, while alluding to the variable recovery outcomes often associated with neurological trauma, offers a potential avenue for both stratifying injury severity and predicting clinical outcomes based on residual serotonin levels in the cord. They also suggest that there may be a residual-serotonin threshold below which physical rehabilitation may only be marginally effective. Serotonergic interventions could substantially improve motor-recovery outcomes in those cases. These observations allude to a potential role for residual serotonin as a neurochemical biomarker of SCI severity. Investigation of serotonin levels in the cerebrospinal fluid (from the spinal cavity), through mass-spectrometry approaches (88, 89), may provide clinicians with critical information that can be used for personalizing treatments wherein serotonergic interventions can be prescribed when spinal levels are below a cut-off, which will need evaluation in future investigations.

### Conclusions

Our results demonstrate that intraspinal delivery of quipazine, caudal to the lesion, can significantly improve motor-recovery outcomes promoted by physical rehabilitation after a chronic and severe contusion of the spinal cord. Notably, delivery of quipazine resulted in substantial behavioral restoration that persisted for weeks after the end of therapy (in the absence of quipazine). The ability of our neuromodulatory intervention to induce sustained benefits underscores its clinical relevance and signals a serotonin-driven reorganization of severely-damaged adult spinal circuitry. Subsequent studies will explore less-invasive intrathecal or epidural routes of neuromodulator administration. Finally, our experiments provide important insights into serotonergic modulation of spinal circuitry and highlight a potential role for serotonin as a neurochemical biomarker of injury severity, which will inform future investigations and therapies targeting chronic SCI.

## Materials and Methods

All animal-handling, behavioral training and surgical procedures were approved by the University of Washington’s Institutional Animal Care and Use Committee, and they conformed to the National Institutes of Health’s *Guide for the Care and Use of Laboratory Animals* (90).

### Subjects

Experiments were performed with 20 adult female Long-Evans rats (procured from Charles River Laboratories). A weight cut-off of 250 g (91), which corresponds to an age cut-off of approximately 90 days (92), was imposed (before the SCI) to be considered an adult animal. Pre-injury weights of rats in the two intervention groups ranged between 266 – 354 g.

### Behavioral training

After allowing several days for acclimation, rats were trained to perform a skilled forelimb reach-and-grasp behavior (46) during their dark (*i.e.,* active) cycle, using operant-conditioning protocols (47) for food-pellet (Bio-Serv, F0299) reward. To initiate training, animals were placed in a custom-designed clear plexiglass arena that contained two vertical slits near the lower-front corners. A food pellet was inserted inside a small dimple in a (custom-made) plastic block that was placed outside the arena in front of one of the vertical slits. A gap between the arena and the plastic block ensured that the food pellet would fall and land out of reach when not firmly grasped by the rats. After identification of the dominant forelimb, rats were trained to reach out through the slit, grasp the food pellet placed on the block with their dominant forepaw, and bring the pellet through the slit into the arena and to their mouth. The width and position of the slit were chosen to ensure that rats could reach the pellets only with one forelimb. After SCI, this design discouraged reaching with the contralesional forelimb, thus focusing retraining on the impaired limb.

### Behavioral scoring

Each behavioral session consisted of 20 pellet-retrieval trials, which were preceded by a short unscored warm-up of five trials. Performance was quantified by the cumulative total of the outcome of individual trials in the session. There were five possible outcomes for a pellet-retrieval attempt: [1] the rat retrieved the food pellet and brought it inside the arena to its mouth with the dominant forelimb (which was counted as a success), [2] the rat retrieved the food pellet, but the pellet dropped inside the arena on the way to its mouth (considered a drop-inside), [3] the rat retrieved the food pellet, but the pellet dropped outside the arena (scored as a drop-outside), [4] the rat extended the dominant forelimb far enough to touch the food pellet, but no grasp was involved (counted as a touch), or [5] the rat made no contact with the pellet (which was considered a miss). These outcomes (success, drop-inside, drop-outside, touch, miss) were quantified based on the level of reach and grasp ability involved in the effort. The rat was assigned a reach score of 0.50 for success, drop-inside, drop-outside and touch. These different outcomes received the same reach score due to the similar level of forelimb extension involved in each of the outcomes. In contrast, the rat received a grasp score of 0.50, 0.33 and 0.17 for success, drop-inside and drop-outside, respectively. A touch received a grasp score of 0 (since the rat did not lift the pellet out of the dimple in the block). These scores were graded based on the grasping ability involved in each outcome. Trials that resulted in a miss received 0 for both reach and grasp. Individual reach and grasp scores were summed up to generate the combined reach-and-grasp score per trial. This scoring scheme is summarized in Table S1. Next, scores from the 20 individual trials were added to compile behavioral scores for the session. Ten sessions of forelimb-motor training occurred per week, with the sessions divided across the five weekdays. Daily sessions were approximately 30 minutes apart to prevent confounds due to fatigue. Scores from the ten sessions were averaged and expressed as a percentage of the 20 trials to compile weekly reach, grasp and reach-and-grasp performances. Training was complete when rats attained a weekly reach-and-grasp performance score of at least 80% (which took 1 – 4 weeks).

### Spinal contusion

After rats attained a reach-and-grasp performance score of at least 80% on the forelimb-motor task, they received a unilateral moderate-to-severe dorsal contusion injury at the border of C4 – C5 spinal segments on their dominant forelimb side (83), using an Ohio State impact device (48). Briefly, rats were anesthetized with a mixture of ketamine (70 mg/kg) and xylazine (10 mg/kg), administered intraperitoneally, and received local intradermal anesthetics (1% lidocaine and 0.25% bupivacaine) along the incision line and subcutaneous antibiotics (enrofloxacin, 0.70 mg/kg) pre-operatively. In an aseptic procedure, the dorsal neck musculature over the cervical spinal column was retracted, and a hemilaminectomy was performed on the C4 vertebrae on the dominant forelimb side. Clamps of the Ohio State device were then placed on the lateral processes of C3 and C5 vertebrae to fix the spinal column into a suspended position. Next, the tip of the impactor probe was aligned at the dorsal root entry zone of the exposed cord, lowered until probe sensors detected contact, and then triggered to rapidly displace the cord (for 14 milliseconds) to simulate an impact injury. Injury displacements of the animals in our study ranged between 0.61 – 0.71 mm. After the contusion, the rat was removed from the clamps, and the musculature and skin were sutured closed (with 4-0 polyglycolic acid and 5-0 nylon suture, respectively; AD Surgical). Rats recovered with fluid and heat support. Injured animals received post-operative drugs (1 mg/kg subcutaneous sustained-release buprenorphine and 0.20 mg/mL enrofloxacin in their water) and manual bladder expressions (two times per day) until voiding reflexes returned, usually within 1 – 4 days.

All animals exhibited near-complete paralysis of the ipsilesional forelimb in the first 1 – 2 days after SCI. Pronounced ipsilesional forelimb deficits persisted chronically and were characterized by a flexion bias at the elbow, wrist and digits, and reduced ability to volitionally extend the limb. Paw placement during exploratory rearing behavior was generally asymmetrical and dominated by the contralesional forelimb, which was also dominant during weight-bearing, grooming and locomotion. Additionally, movements of the impaired forelimb were generally slower and lacked precise coordination. In the severely-injured animals, spontaneous recovery was negligible, and near-complete paralysis of the ipsilesional forelimb remained present at the start of the intervention.

### Intraspinal cannula and subcutaneous pump implants

Four weeks after SCI, rats were implanted with a custom-designed cannula whose distal tip terminated in the ipsilesional C7 spinal segment (caudal to the injury site; see experimental timeline in Figure 1A). The cannula assembly consisted of a 3-mm piece of 0.13-mm inner-diameter polyurethane spinal catheter (Instech Laboratories, BTPU-010) of which 2 mm were inserted into (180 mm of) an intermediate-sized silicone tubing (whose inner diameter was 0.30 mm; DuPont, 508-001). The overlap junction was secured with 4-0 silk suture (AD Surgical) and medical-grade liquid silicone adhesive (Factor II). Next, a custom-cut polyamide disc was placed at the junction to act as a (subdural) depth stop for limiting the penetration depth to 1 mm. The intermediate tubing served to interface the spinal catheter with the larger-diameter outlet tubing from a peristaltic pump (Alzet, iPrecio SMP-200; 50), which was implanted subcutaneously in the same surgery or in a separate procedure (as detailed below) and was used for controlled delivery of fluids into the cord.

Rats were anesthetized using inhaled isoflurane (2 – 3% in oxygen), placed on a heating pad, and administered subcutaneous antibiotics (enrofloxacin, 0.70 mg/kg), corticosteroids (dexamethasone, 0.50 mg/kg), analgesics (meloxicam, 1 mg/kg) and lactated Ringer’s solution (for fluid support) pre-operatively. Surgical plane was verified *via* toe pinch and ocular reflex, and vitals were monitored continuously throughout the surgery using a SomnoSuite surgical monitoring system (Kent Scientific). Under aseptic conditions, an incision was made along the dorsal midline above C7, the muscle was separated and retracted, and the spinal column was exposed. A full C7 and a partial-to-full C6 hemilaminectomy was performed on the ipsilesional vertebrae. The cannula was flushed with 70% isopropanol and sterile saline, then primed with saline, and tied shut at the proximal (or the silicone) end using 4-0 silk suture. Next, the dura mater over the C7 dorsal root entry zone was incised and the polyurethane end of the cannula was inserted 1 mm into the cord, as set by the depth stop. The dura mater was then closed atop the depth stop and around the cannula with 7-0 silk suture (AD Surgical) to anchor it in place. The remaining length of the cannula extended dorsally through the muscle (which was closed with 4-0 polyglycolic acid suture) and was coiled inside a subdermal pocket.

During the cannula-implant surgery or in a separate brief procedure (that occurred at the end of the pre-intervention week and before the start of therapy), a second incision was made several centimeters rostral to the cannula-implant site and a subdermal pocket was created *via* blunt dissection for placing a battery-operated microinfusion pump (Alzet, iPrecio SMP-200; 50) whose delivery profile was programmed before implantation. The pump-outlet tubing was trimmed to 20 mm, and the reservoir and outlet tubing were primed with saline. Next, the cannula was tunneled subdermally to the pump pocket, 5 mm of its proximal length (at the silicone end) were snipped, and it was inserted inside the pump-outlet tubing. The junction was secured using 4-0 silk suture and liquid silicone (World Precision Instruments, Kwik-Sil). Note that an additional small incision was made above the cannula-implant site to expose the proximal (or silicone) end of the cannula when the pump was implanted in a separate procedure. Lastly, the pump was anchored inside the subdermal pocket using silicone-coated 4-0 silk suture, and the incision sites were sutured closed (with 5-0 nylon). Rats recovered on a heating pad and were administered analgesics (1 mg/kg subcutaneous sustained-release buprenorphine) and antibiotics (0.20 mg/mL enrofloxacin in their water). Animals were allowed two weeks of post-operative recovery before assessment of their injury severity.

### Impairment matching and group assignments

Saline was delivered during two weeks of post-operative recovery, which was followed by pre-therapy motor assessment on the previously-mastered pellet-retrieval task (46) during week 7 (also in the presence of saline; see experimental timeline in Figure 1A) to quantify the functional deficits produced by the cervical contusion. To account for the variability in motor impairment across animals arising from the large range of injury severities tested in our study, we used an impairment-matched design, which involved classifying rats into different injury-severity levels based on their pre-therapy reach-and-grasp performance scores. Within a given injury-severity level, rats were then randomly assigned to one of the two intervention groups. Table S2 details this classification, showing the minimum, maximum and average pre-therapy motor performances of rats tested at each injury-severity level.

To assess the effect of therapy on forelimb-motor recovery as a function of the severity of the cervical injury, rats with a pre-therapy reach-and-grasp motor performance score (normalized to pre-injury levels) of less than or equal to 30% were ranked severely injured (Table S2, injury-severity levels I_4_ and I_5_) and rats with a pre-therapy performance of greater than 30% were classified as moderately injured (Table S2, injury-severity levels I_1_ – I_3_) in our study.

### Therapy and post-therapy follow-up

Before the start of therapy, animals were briefly sedated using inhaled isoflurane, and the fluid in the pump reservoir was exchanged from saline to quipazine (Sigma-Aldrich, Q1004), *via* a percutaneously-accessible septum, in rats that received the combined intervention. To maintain uniformity of procedures across all animals, a saline-to-saline exchange was performed in the control animals.

Therapy, consisting of physical rehabilitation and neuromodulator or vehicle delivery, was administered during weeks 8 – 13 after SCI (see experimental timeline in Figure 1A). The eight-week delay between injury and start-of-therapy was chosen to allow the contusion to translate to a chronic SCI model (51, 52). Quipazine/vehicle was delivered intraspinally below the lesion for approximately 4.5 hours per day (between 11:00 – 15:24) for six weeks. Pumps were pre-programmed (before implantation) to gradually increase the flow rate from 0.5 – 4.0 µL/hour during the therapy window, with the flow rate doubled every 30 minutes and then maintained at 4.0 µL/hour. A gradual delivery profile was chosen due to its association with more durable effects, such as sustained activation of receptors and downstream signaling pathways, compared to fast chemical application (93). Pumps stayed off during the remaining hours of the day (*i.e.,* outside the therapy window). Rats received 20 µg/week of quipazine. The therapy duration of six weeks was chosen due to the plateau in gains observed at this time point (Figure 1B). Physical rehabilitation consisted of ten sessions per week of retraining of the impaired forelimb on the pellet-retrieval task, divided between the five weekdays, and occurred during periods of ongoing neuromodulator/vehicle delivery. Each physical-training session, in turn, consisted of 20 pellet-retrieval trials, preceded by a short warm-up (as described above in this section).

After completion of therapy, a reverse fluid exchange from quipazine to saline was performed under brief isoflurane anesthesia in rats in the combined intervention group, while a (second) saline-to-saline exchange was performed in the control animals. Rats continued to be tested in the pellet-retrieval task for an additional four weeks (in the presence of saline; weeks 14 – 17; see experimental timeline in Figure 1A) to assess if neuromodulator-mediated gains persisted in the absence of quipazine. Note that due to the pre-programmed nature of the pumps, they could not be turned on or off on-the-fly. Moreover, constraints arising from the limited number of delivery steps available in the programming software precluded different weekly infusion profiles. Therefore, saline was delivered between 11:00 – 15:24, using the same routine detailed above, during the follow-up period (rather than keeping pumps off).

Pumps were refilled with quipazine/saline as needed during therapy and follow-up periods. Reservoir-volume checks on the pumps were performed during all exchanges/refills to confirm fluid delivery and rule out any clogging issues. Clogging was observed in two rats in the control group who did not receive the entire volume of saline. Due to the long-term nature of the experiments and their assignment within the control group, these rats were included in the study.

All rats in both intervention groups received the same amount of retraining on pellet retrieval during (per week) pre-therapy assessment, therapy and follow-up periods. The pellet-retrieval task served as both a form of use-dependent physical training and a means to quantify changes in forelimb-motor performance. To eliminate biases in behavioral scoring, all retraining was performed by researchers blinded to the group assignments of the rats. Additionally, these individuals were not involved in data analyses or interpretation. Moreover, surgeries and procedures were similar in all animals, and personnel involved in surgeries and procedures, related to fluid exchanges and reservoir refills, did not train animals.

### Terminal surgery

After completion of post-therapy follow-up, a terminal surgery was performed in all rats from both intervention groups, during week 18 (see experimental timeline in Figure 1A), in which we ensured that pumps were still operational at the end of the study. We also verified the integrity of fluidic junctions and flow through the entire line. Next, we checked the location of the distal tip and entrance angle of the cannula to confirm that fluid was delivered in the spinal cord (*vs.* in the spinal cavity or epidurally) caudal to the lesion. Lastly, we looked for signs of infection in the spinal cord and muscle layers. No infection was seen in any of the rats in the study.

### Behavioral analyses

Forelimb-motor performance on the pellet-retrieval task was the primary outcome measure employed in our study to assess both the chronic functional deficits produced by the cervical-contusion injury and intervention-mediated changes in motor function. We quantified the reach and grasp motor performance, individually and together, after SCI relative to pre-injury performance levels. The weekly motor performance during the pre-therapy, therapy and follow-up periods was expressed as a percentage of the rat’s individual pre-SCI maximum, which was the maximum weekly score obtained during behavioral training. Additional details can be found in SI Materials and Methods. Note that, while the ability to reach could be assessed independently, the assessment of grasp was dependent on the rat accurately extending its forelimb outside the arena far enough to reach the food pellet, implying that only pellet-retrieval trials in which rats achieved a touch, drop-outside, drop-inside or success could be used for evaluating grasp. While this could potentially result in an underestimation of grasp ability (due to its dependence on reach), there were only two (out of 40) sessions during the four-week post-therapy follow-up in one control rat alone (out of the 20 rats in our study across the two intervention groups) in which the rat was unable to reach the pellet and all trials resulted in misses. These two sessions were, therefore, excluded from the grasp analysis.

We also calculated the therapeutic gains, which was the difference between the (mean) post-therapy and pre-therapy motor performances, produced by our interventions. The post-therapy performance was calculated by averaging scores obtained during weeks 14 – 17.

### Statistics

Motor-performance scores and therapeutic gains are reported as mean ± standard error (unless otherwise stated). Paired-samples *t* tests were employed for comparisons of performance scores/gains between the two impairment-matched intervention groups (53, 54) and across (two) time points (such as in Figure 2). Before performing paired *t* tests, we verified that the differences between the paired data followed a normal distribution. Normality was assessed using the Shapiro-Wilk test. Wilcoxon signed-rank test, which is the non-parametric equivalent of the paired-samples *t* test, was used when the data were not normally distributed. We also used the Friedman test, which is the non-parametric equivalent of repeated-measures analysis of variance, to compare data when more than two time points were involved. All statistical analyses were performed in IBM SPSS Statistics (version 27). Significance for all statistics was set at the .05 level.

In addition to assessing statistical significance, we quantified the magnitude of the effect using the “effect size” metric, which is the standardized difference between the means of two groups (94). Additional details of effect-size calculations can be found in SI Materials and Methods. Effect-size descriptors developed by Cohen (94) and extended by Rosenthal (95), which are summarized in Table S3, were used for qualitative interpretation of the magnitude of observed effects.

## Acknowledgements

We thank Gawin Mai, Pierce Lovinger, Juliana Baranov and Amulya Jolepalem for behavioral training of rats and assistance with handling and care of the animals. This work was supported by grants from the Craig H. Neilsen Foundation (384977), the National Institutes of Health (NS099872), the Paralyzed Veterans of America (3082) and the University of Washington Royalty Research Fund (A166877). Additionally, the study was supported by an Early Investigator Catalyst Award from the Institute of Translational Health Sciences at the University of Washington.

