## Supporting Information for "Serotonergic Modulation of Spinal Circuitry Restores Motor Function after Chronic Spinal Cord Injury"

This file includes the following:

SI Materials and Methods (page 2)

Supplementary Tables (pages 3 – 5)

SI References (page 6)

List of Supplementary Tables:

Table S1 Behavioral scoring system

Table S2 Impairment-matched intervention groups

Table S3 Thresholds for interpreting effect sizes

### SI Materials and Methods

**Behavioral analyses.** To assess changes in forelimb-motor performance, the average weekly reach-and-grasp score after SCI was expressed as a percentage of the rat's individual pre-SCI maximum, which was the maximum weekly score obtained during behavioral training, according to the following equation:

$$\text{Reach-and-Grasp Performance (\%)} = \frac{\text{Weekly Post-SCI Reach-and-Grasp Score}}{\text{Maximum Pre-SCI Reach-and-Grasp Score}} \times 100$$

Similarly, to evaluate changes in the reach and grasp performances, individually, we expressed the (average) weekly reach or grasp score after SCI as a percentage of the rat's individual pre-SCI maximum, according to the following equations:

$$\text{Reach Performance (\%)} = \frac{\text{Weekly Post-SCI Reach Score}}{\text{Maximum Pre-SCI Reach Score}} \times 100$$

$$\text{Grasp Performance (\%)} = \frac{\text{Weekly Post-SCI Grasp Score}}{\text{Maximum Pre-SCI Grasp Score}} \times 100$$

Note that normalized motor performances, calculated using the above equations, that exceeded 100% (due to the post-SCI score being greater than the rat's pre-SCI maximum) were scaled back to 100%.

**Statistics.** In addition to assessing statistical significance, details of which are provided in Materials and Methods (in the Main Text), we quantified the magnitude of the effect, when significant, using the "effect size" metric (1), which is the standardized difference between the means of two groups, according to the following equation:

$$\text{Effect Size} = \frac{(\text{Mean of Experimental Group} - \text{Mean of Control Group})}{\text{Population Standard Deviation}}$$

Population standard deviation was estimated by pooling the (sample-size-weighted) standard deviations of the experimental and control groups, using the equation below, when standard deviations of the two groups were similar.

$$\text{Pooled Standard Deviation} = \text{SQRT}\left(\frac{(N_E - 1) * SD_E^2 + (N_C - 1) * SD_C^2}{N_E + N_C - 2}\right)$$

Here, $N_E$, $SD_E$, $N_C$ and $SD_C$ are the sample sizes and standard deviations of the experimental and control groups, respectively. To assess similarity between standard deviations, we calculated the difference between the coefficient of variation (which is the ratio of the standard deviation to the mean) of the two groups and imposed a cut-off of 5%. When this difference exceeded 5%, the standard deviations of the two groups were considered to not be similar in which case the standard deviation of the control group alone was used as our population standard deviation. These metrics, when estimated using the pooled standard deviation *vs.* the standard deviation of the control group, correspond to Hedges' g (2) and Glass' delta (3) effect-size indices, respectively. Lastly, effect-size descriptors developed by Cohen (1) and extended by Rosenthal (4), which are summarized in Table S3, were used for qualitative interpretation of the magnitude of observed effects.

### Supplementary Tables

**Table S1.** Behavioral scoring system.

| SCORE | Success | Drop-Inside | Drop-Outside | Touch | Miss |
|---|---|---|---|---|---|
| Reach | 0.50 | 0.50 | 0.50 | 0.50 | 0 |
| Grasp | 0.50 | 0.33 | 0.17 | 0 | 0 |
| Reach-and-Grasp | 1 | 0.83 | 0.67 | 0.50 | 0 |

Depending on the outcome of a pellet-retrieval trial, rats received differential reach and grasp behavioral scores, which were summed up to generate the combined reach-and-grasp score for the trial.

**Table S2.** Impairment-matched intervention groups.

| INJURY-SEVERITY LEVEL | Minimum Pre-Therapy Performance | Maximum Pre-Therapy Performance | Mean Pre-Therapy Performance for Physical Training Alone | Mean Pre-Therapy Performance for Physical Training & Quipazine | Number of Rats per Intervention Group |
|---|---|---|---|---|---|
| $70 < I_1 \leq 80$ | 70.4 | 78.0 | 73.34 ± 4.21 | 76.85 ± 1.68 | 2 |
| $50 < I_2 \leq 70$ | 55.9 | 66.8 | 66.76 | 55.89 | 1 |
| $30 < I_3 \leq 50$ | 35.7 | 47.5 | 37.25 ± 2.22 | 42.54 ± 7.06 | 2 |
| $15 < I_4 \leq 30$ | 16.1 | 30.0 | 24.90 ± 5.13 | 17.82 ± 1.47 | 3 |
| $5 < I_5 \leq 15$ | 6.7 | 11.2 | 9.37 ± 2.64 | 7.90 ± 1.71 | 2 |

Rats were classified into five injury-severity levels, $I_1 – I_5$, based on their pre-therapy reach-and-grasp motor performance, with rats in $I_1$ being the least injured and rats in $I_5$ being the most injured. Columns 2 and 3 show the minimum and maximum performance values (respectively) at the five injury-severity levels across the two intervention groups. Columns 4 and 5 show the pre-therapy performance means and standard deviations for rats that received saline and quipazine, respectively. The reach-and-grasp performance of rats is expressed as a percentage of their individual pre-SCI maximum score. The last column shows the number of rats that were tested per intervention at each level of injury severity.

**Table S3.** Thresholds for interpreting effect sizes.

| EFFECT SIZE | Interpretation |
|---|---|
| $0.00 < d < 0.20$ | Very Small or Trivial |
| $0.20 \leq d < 0.50$ | Small |
| $0.50 \leq d < 0.80$ | Medium |
| $0.80 \leq d < 1.30$ | Large |
| $d \geq 1.30$ | Very Large |

The rationale for these benchmarks was outlined by Cohen (1). Supplementing Cohen's definition of small, medium and large effect sizes, Rosenthal extended the classification to very large effects (4). *d*, standardized mean difference (or effect size).
